# Mathematical analysis of light-sensitivity related challenges in assessment of the intrinsic period of the human circadian pacemaker

**DOI:** 10.1101/2023.07.14.549062

**Authors:** Imran M. Usmani, Derk-Jan Dijk, Anne C. Skeldon

## Abstract

Accurate assessment of the intrinsic period of the human circadian pacemaker is essential for a quantitative understanding of how our circadian rhythms are synchronised to exposure to natural and man-made light-dark cycles. The gold standard method for assessing intrinsic period in humans is forced desynchrony (FD) which assumes that the confounding effect of light on assessment of intrinsic period is removed by scheduling sleep-wake and associated dim light-dark (LD) cycles to periods outside the range of entrainment of the circadian pacemaker. However, the observation that the mean period of free-running blind people is longer than the mean period of sighted people assessed by FD (24.50 *±* 0.17 h versus 24.15 *±* 0.20 h, p *<* 0.001) appears inconsistent with this assertion. Here, we present a mathematical analysis using a simple parametric model of the circadian pacemaker with a sinusoidal velocity response curve (VRC) describing the effect of light on the speed of the oscillator. The analysis shows that the shorter period in FD may be explained by exquisite sensitivity of the human circadian pacemaker to low light intensities and a VRC with a larger advance region than delay region. The main implication of this analysis, which generates new and testable predictions, is that current quantitative models for predicting how light exposure affects entrainment of the human circadian system may not accurately capture the effect of dim light. The mathematical analysis generates new predictions which can be tested in laboratory experiments. These findings have implications for managing healthy entrainment of human circadian clocks in societies with abundant access to light sources with powerful biological effects.

## Introduction

Appropriate timing of physiology and behaviour to temporal niches associated with geophysical cycles contributes to fitness of biological systems (West and Bechtold, 2015), including the health of humans (Fishbein et al., 2021). This ‘appropriate timing’ is reflected in 24-hour rhythmic variation in gene expression, translation, physiology and behaviour and is referred to as circadian rhythmicity. A defining feature of circadian rhythms is that they are self-sustaining (Pittendrigh, 1960). The rhythms are generated by oscillators, whose activity persists in the absence of cyclical changes, known as zeitgebers, in the external environment (Aschoff, 1960). In the study of the central circadian pacemaker in mammals, the intrinsic period refers to the period of the pacemaker in the absence of zeitgebers. The intrinsic period is close to, but rarely equal to, 24 hours (Czeisler et al., 1999) and entrainment to 24 hours is achieved by an adjustment to the intrinsic rhythm of the pacemaker through exposure to 24-hour zeitgebers (Daan, 1977, 2000).

Accurate estimation of the intrinsic period is important for two main reasons. First, the intrinsic period is a key factor in determining whether the pacemaker can entrain to 24-hour light-dark (LD) cycles, since the magnitude of the adjustment required for entrainment depends on the difference between the intrinsic period and the period of the LD cycle (Pittendrigh and Daan, 1976b). Second, when the pacemaker entrains to LD cycles, the intrinsic period determines the phase of entrainment, i.e., the timing of endogenous rhythmicity relative to the zeitgeber, with longer intrinsic period associated with later sleep timing (Duffy et al., 2001). Thus the intrinsic period of the circadian pacemaker and its variation between individuals informs the interpretation of circadian rhythm sleep-wake disorders (Meyer et al., 2022; Micic et al., 2016) as well as the variation in the timing of rhythmicity in the general population. For example, mathematical models suggest that those with a longer intrinsic period are more sensitive to the delaying effects of access to evening light (Skeldon et al., 2017). Furthermore, guidelines on healthy light exposure requirements critically depend on an assessment of the average and between individual variation of this key parameter along with the sensitivity of the pacemaker to light.

The most reliable way to assess the intrinsic period of the pacemaker is to place a person or animal in DD. This is because in constant darkness (DD), the principal zeitgeber to the pacemaker, namely light (Czeisler et al., 1981; Dijk et al., 1995), is removed. The rest-activity cycle and behaviours associated with the rest-activity cycle, such as feeding, persist in DD, but the non-uniform distribution of these behaviours across the circadian cycle are assumed not to affect the period of the pacemaker to a significant extent, but see (Kas and Edgar, 2001). In nocturnal animals, the period in DD is measured readily (Pittendrigh and Daan, 1976a), but there are practical and ethical barriers to studying sighted humans in DD. Consequently, sighted people are rarely studied in DD, although in the 1970’s Rütger Wever did assess the intrinsic period of five sighted humans who lived in DD for approximately two weeks (Wever, 1979). The intrinsic period of humans has traditionally been assessed in classical free-run (Wever, 1979) and forced desynchrony (FD) protocols (Czeisler et al., 1999; Wang et al., 2023). In FD, participants are exposed to LD cycles with a period very different from 24 h, usually 28 h or 20 h. In standard protocols, lights are on and participants are required to be awake for two-thirds of the time. Lights are off and participants are in bed and encouraged to sleep for the remaining one-third. With 28 (20) h cycles, wake is therefore scheduled to occur 4 h later (earlier) each day. Since 28 (20) h is outside the limits of entrainment, over the course of (an integer multiple of) 6 light / dark cycles the circadian clock is exposed to light at (approximately) all different phases. In this design, the aim is to minimise the effects of light, to allow the human circadian clock to progress at its natural period.

More recently, the intrinsic period has been assessed by measuring the *in vitro* period of fibroblasts taken from individual participants (Pagani et al., 2010). The fibroblast period is measured by introducing firefly luciferase genes into the fibroblast cells via a lentivirus. Since the expression of firefly luciferase is then driven by the circadian gene Bmal1, the fibroblasts exhibit periodic patterns of bioluminescence which are measured via luminometry.

Fig. 1 summarises estimates of the intrinsic period in blind and sighted humans using these various methods. Here, only blind participants with non-entrained rhythms are included, where blind means having no subjective perception of light. For studies where melatonin suppression was measured, we have further restricted to those who had no melatonin suppression by light. For example, FlynnEvans et al. (2014) studied 127 blind people of whom 41 had no light perception and 16 of these were nonentrained. In Fig. 1, only the 16 nonentrained participants are included.

**Figure 1:**
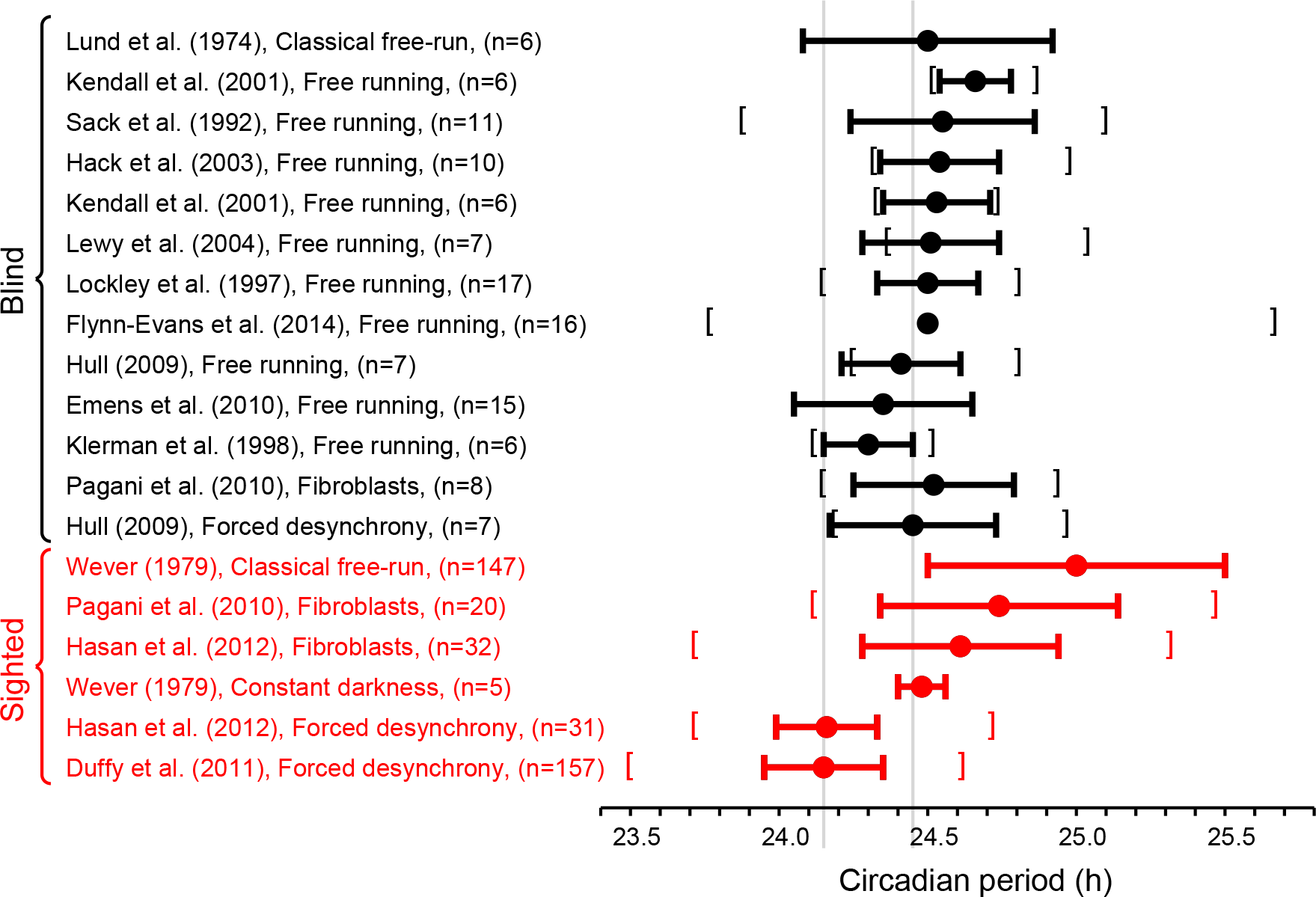
Estimates of the intrinsic period of the circadian pacemaker in blind and sighted humans using various protocols. Circles indicate the mean with the horizontal bars indicating the mean +*/−* the standard deviation. Where available, the square brackets indicate the range of the measurements. In some cases (e.g. Lewy et al., 2004) where the studies only include a small number of participants the distribution of periods is skewed so that the smallest value recorded is greater than the (mean*−*standard deviation). The grey vertical lines indicate accepted values for the circadian period of sighted (as measured in forced desynchrony) and blind individuals respectively.

In Fig. 1 it can be seen that the mean period of blind people is consistent across different protocols (Emens et al., 2010; Hack et al., 2003; Hull, 2009; Kendall et al., 2001; Klerman et al., 1998; Lewy et al., 2004; Lockley et al., 1997; Lund et al., 1974, Sack et al., 1992), and consistent within individuals assessed in both free-running field conditions and in an FD protocol in the laboratory (Hull, 2009). In contrast, the mean period of sighted people is variable depending on the protocol (Duffy et al., 2011; Hasan et al., 2012; Wever, 1979). The period of sighted people in DD is consistent with intrinsic period estimates in the blind. In addition, there is no significant difference between the mean period of fibroblasts from sighted people and the period of fibroblasts from blind people (p = 0.17). The reported standard deviations of estimates of periods appear larger in classical free run and fibroblasts than observed in FD protocols but are similar between sighted people in FD and assessments in the blind.

Motivated by concerns over the impact of room lighting in classical free run protocols, forced desynchrony (FD) has emerged as a widely accepted gold standard method for assessing the intrinsic period of the circadian pacemaker in sighted humans (Dijk et al., 2020; Wang et al., 2023). It has been proposed that the shorter period of sighted people in FD compared to blind people is due to after-effects of prior entrainment in sighted people (Duffy et al., 2005). The presence of after-effects implies that the period of sighted people in FD should be variable depending on prior period of entrainment. However, in humans, the period of the zeitgeber during prior entrainment appears to have only a modest effect on the subsequent period of the pacemaker in FD (mean difference 0.1 h) (Scheer et al., 2007). It is also interesting, and maybe surprising, to note that the average period of fibroblasts in both sighted and blind people *in vitro* are comparable with each other and similar to the intrinsic period of blind people, see Fig. 1, although in sighted participants the fibroblast period does not correlate with the period of plasma melatonin as assessed in FD (Hasan et al., 2012).

In classical free-run, the self-selected light exposure of participants is likely to modulate the period of the pacemaker in sighted people (Klerman et al., 1996). In order to minimise the effect of light, FD protocols aim to distribute light evenly over the circadian cycle and use dim light. For example, in Wang et al. (2023) it is recommended that light levels in FD should be less than 15 lux, and it is reported that in many FD experiments light of intensity less than four lux has been used. Even at these low intensities, there is evidence that dim LD cycles may modulate the period of sighted people. For example, Wright et al. (2001) demonstrated that sighted people can entrain to dim (*≈* 1.5 lux) 24-h LD cycles in a carefully controlled experiment with an imposed 8:16 rest:activity cycle.

In view of these discrepancies and unresolved issues relating to the intrinsic period of the human circadian pacemaker, and an absence of a formal mathematical analysis of how light may affect the human circadian pacemaker, a further analysis seems warranted. Here, we use a simple mathematical model of the circadian pacemaker to describe the effect of dim LD cycles on the circadian pacemaker in sighted humans. Using this model, we derive an expression relating the period in FD to the intrinsic period, which highlights the dependence of assessed period on the symmetry of the velocity response curve in the model. We estimate parameters of our model using Wright’s data on the entrainment of humans to dim LD cycles (Wright et al., 2001). Then, we present a hypothesis for the observed shorter period of sighted people in FD compared to blind people. We describe experimental protocols to test this hypothesis. Our hypothesis offers one solution to a long-standing discrepancy and has implications for quantitative models that predict the effect of the light environment as mandated by policies about light exposure requirements and work schedules.

## Methods

### Simple clock model of the human circadian pacemaker for dim light conditions

It is well established that the human circadian pacemaker behaves as a phase-amplitude oscillator, perturbations of which can lead to changes in phase and amplitude (Czeisler et al., 1989; Khalsa et al., 1997; Strogatz, 1990). Kronauer’s model of the human circadian pacemaker (Jewett and Kronauer, 1998) and its later versions (Forger et al., 1999; Jewett et al., 1999; St Hilaire et al., 2007) are the most widely used in human circadian research. These models were designed to replicate phase resetting studies, including amplitude reduction, type-1 and type-0 phase resetting (Khalsa et al., 1997). These models are currently being used to predict human circadian phase from ambulatory light data (Huang et al., 2021; Rea et al., 2020; Woelders et al., 2017) across a range of populations including students (Phillips et al., 2017) and shiftworkers (Stone et al., 2019) and are competitive with traditional phase assessment methods in terms of accuracy (Dijk and Duffy, 2020). Kronauer-type models have been used to suggest interventions to minimise the disruptive effects of jet-lag (Serkh and Forger, 2014), non-24 h sleep/wake disorder, shiftwork and social jetlag (Diekman and Bose, 2022). Kronauer-type models have also been combined with models of sleep regulation to investigate changes in sleep timing preferences (Phillips et al., 2010; Skeldon et al., 2016), sleepiness and cognitive performance due to shift-working (Postnova et al., 2014; Postnova et al., 2018), the impact of light and social constraints on sleep timing preferences and social jet lag (Skeldon et al., 2017), the effects of daylight saving (Skeldon and Dijk, 2019), and used to propose quantitative light ‘availability’ interventions (Skeldon et al., 2022) to normalize sleep timing.

All the Kronauer-type models use a two-dimensional oscillator with a strongly attracting limit cycle to reproduce the self-sustaining activity of the clock coupled with a model for the effect of light on the clock. Earlier versions of the model were developed from experiments in which the light intensity varied from 10 lux to 9500 lux. The most recent version (St Hilaire, 2007) was adapted to more accurately reflect light sensitivity for intensities below 150 lux and additionally includes a non-photic zeitgeber. Here, the focus is on the effect of dim light, which constitutes a weak zeitgeber. It has been established that models with a strongly attracting limit cycle when exposed to a weak zeitgeber are well-approximated by phase only models (Guckenheimer and Holmes, 1983) (see the Supplementary Material for further details). Therefore, for the purpose of analyzing the effects of dim light, here we use a simple phase-only model. In addition, we assume that the effect of light is to continuously modulate the velocity of the clock, which means that the model is parametric. Parametric models are generally considered to be good models of the circadian system in diurnal animals (Daan, 2000).

In phase-only models, the state of the clock at any time is described only by its phase *ϕ ∈* 𝕊^1^. In line with experimental conventions, we specify that *ϕ* = 0 represents the *circadian minimum*, which is the state of the pacemaker when the core body temperature is at its minimum (CBT_min_). Dim light melatonin onset (DLMO) usually occurs about 7 h before the circadian minimum (Benloucif et al., 2005; Brown et al., 1997; Dijk et al., 1999). With the assumption that the clock velocity is approximately equal to 2*π* (24 h)^*−*1^ between DLMO and CBT_min_, the phase of the pacemaker at DLMO is then *ϕ* =17 *×* 2*π*/24. (The assumption that in dim light and / or darkness the clock progresses approximately uniformly is made in both phase-only models (see equation (1) and implicitly in Kronauer-type models (see the Supplementary Material)) .

The velocity 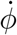 of the clock is the rate of change of phase. In phase-only parametric models,

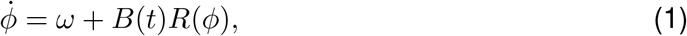

where *ω >* 0 is the intrinsic velocity, *B*(*t*) is the stimulus produced by the LD cycles, *R*(*ϕ*) is the velocity response curve (VRC) to light, and *ϕ ∈ 𝕊*^1^. The intrinsic period of the clock is *τ* = 2*π/ω*. The product *B*(*t*)*R*(*ϕ*) is called the velocity response of the clock to light.

In parametric models, LD cycles of period *T* in which lights are on for *M* of the time produce a stimulus of the form

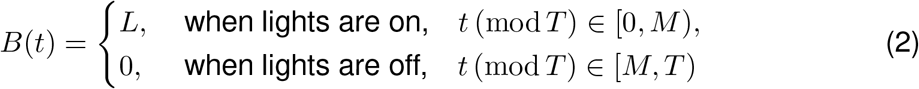

where *L >* 0 and depends on the intensity of light. Here we assume that *L* is constant throughout the light period, which is a reasonable assumption for laboratory protocols. The stimulus in equation (2) represents the *tonic* effect of light on the pacemaker (Daan, 1977).

In parametric models, the VRC typically contains an advance region, in which the effect of a stimulus is to speed up the clock, and a delay region, in which the effect of a stimulus is to slow down the clock. The VRC is a periodic function with period 2*π* so that it can be represented as a Fourier series,

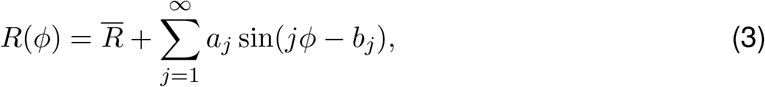

where

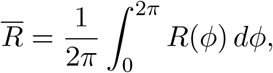

is the mean level of the VRC.

In phase-only parametric models, the effect of light on the velocity of the clock is assumed to be smaller than the intrinsic velocity, that is

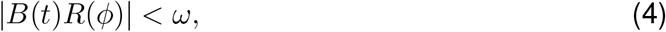

so that the phase of the clock advances monotonically. A schematic of the core body temperature rhythm and an example VRC are shown in Fig. 2.

**Figure 2:**
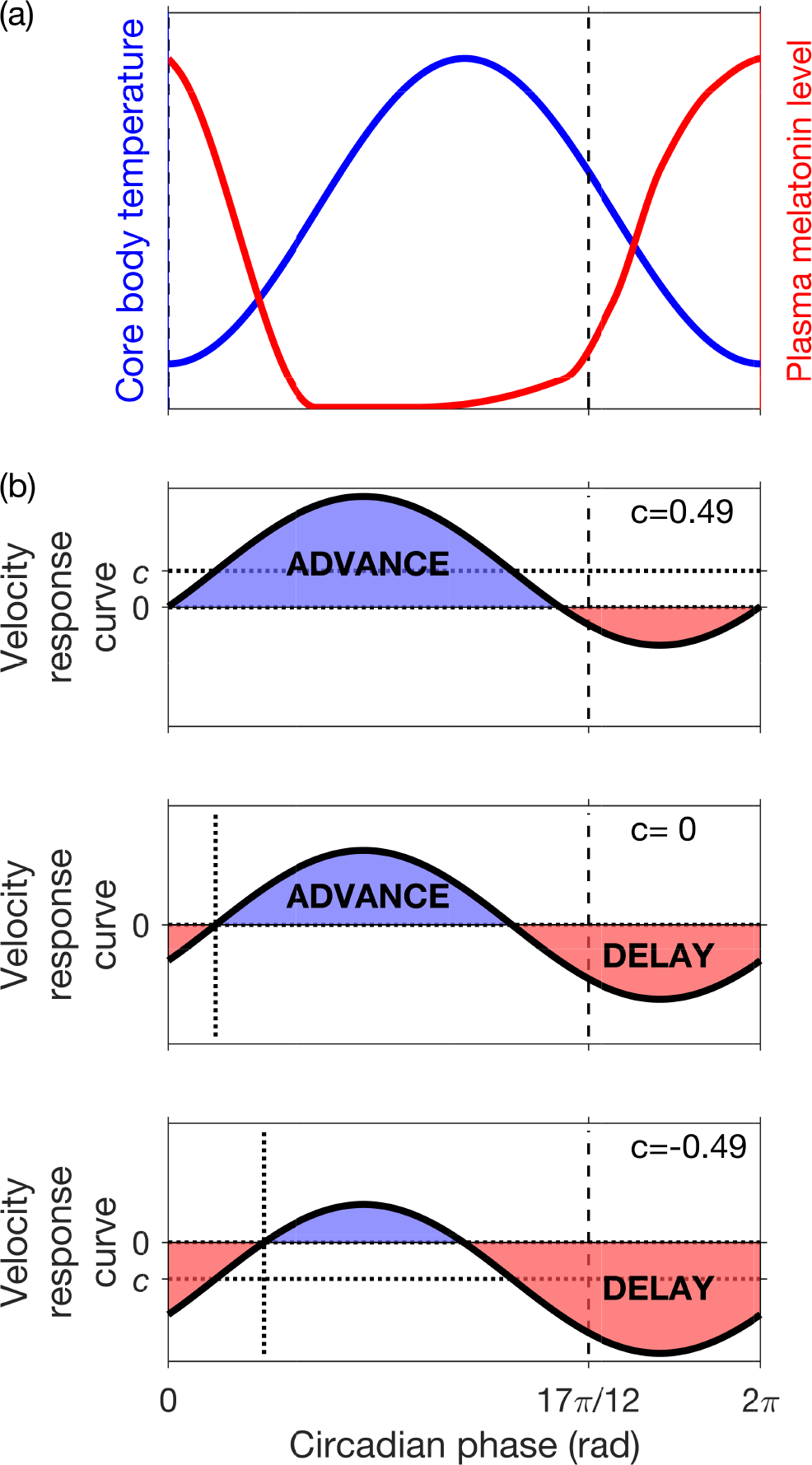
Schematic diagrams of the core body temperature (CBT), plasma melatonin rhythms and the velocity response curve (VRC) in humans. CBT and plasma melatonin rhythms are shown in (a). The minimum of the CBT occurs at circadian phase *ϕ* = 0, and dim light melatonin onset occurs at phase *ϕ* = 17*π/*12. In (b) sinusoidal velocity response curves *R*_1_(*ϕ*) = *c* + sin(*ϕ−b*) are shown for *b* = 0.5 and three different values of the parameter *c*. In each case the VRC has an advance region and a delay region. The sizes and positions of the advance and delay regions depend on the parameters *b* and *c*. The crossover from delay to advance occurs at phase *b−*sin^*−*1^(*c*)*≈b−c* and the width of the delay region is *π−*2 sin^*−*1^(*c*)*≈π−*2*c*, where the approximations are valid for small *c*.

## Results

### Analytical expression for the period in forced desynchrony protocols as found using the simple clock model

In experiments, the mean period in FD is evaluated in one of two ways. When data of a phase marker such as core body temperature is collected throughout the protocol the non-orthogonal spectral analysis (NOSA) algorithm is used. The NOSA algorithm fits a mathematical function consisting of Fourier components for the mean period in forced desynchrony, *τ*_FD_, Fourier components for evoked behaviour with a period of the LD cycle and a correlated noise term (Brown et al., 1997; Czeisler et al., 1999). Alternatively, the interval of time *t* between two occurrences of a biological phase marker such as the CBT_min_ or DLMO is measured. The first occurrence is near the start of the FD protocol, and the second occurrence is near its end. The mean period is given by the quotient *t/n*, where *n* is the number of circadian clock cycles in the interval *t* (Eastman et al., 2015; Scheer et al., 2007).

Using the simple clock model the mean period may be calculated as follows. If the FD protocol consists of *N* LD cycles each with period *T* and the phase of the clock at the start of the protocol is *ϕ*_0_, then the phase at the end of the protocol is found by integrating equation (1) with initial condition *ϕ*(0) = *ϕ*_0_ to find the phase at the end of the protocol, *ϕ*(*NT*),

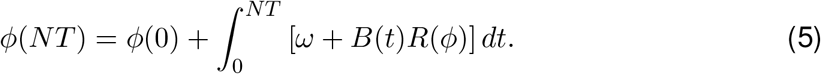

The total change in phase Δ*ϕ*_Tot_ during the protocol is then given by

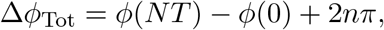

where *n* is the number of times the clock traverses *ϕ* = 0 in the interval *NT* . The mean angular velocity of the clock is

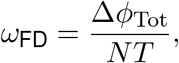

and the mean period of the clock evaluated in FD is

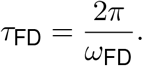

Since the simple clock model does not include noise or evoked effects (e.g. the effect of the sleep-wake cycle on the circadian-regulated core body temperature) and there is no experimental error in calculating the phase, calculating the change in phase from the beginning to end of the protocol should lead to an accurate determination of *τ*_FD_.

In general, it is difficult to carry out the integral in equation (5), and hence find *τ*_FD_, without resorting to numerical methods and simulation. While simulation is an extremely useful technique and has previously been used to optimise FD protocol design (e.g. Lok et al., 2022; Stack et al., 2017), an analytical expression is even more powerful, giving a general understanding of which factors are important. Here, we derive an analytical expression for the period found in FD valid for dim LD cycles and intrinsic periods close to 24 h. We have included (most of) the derivation of the analytical expression in the next two subsections with some steps relegated to the Supplementary Material. The less mathematically-inclined reader may want to skip ahead to equation (14), which gives our expression for the mean period in FD (*τ*_FD_), and the subsequent discussion of the implications and accuracy of our expression.

### Analytical expressions for the phase transition curve and the mean angular velocity

We first construct an expression for the mean angular velocity, *ω*_FD_ in terms of the cumulative phase response across the whole protocol by considering the phase transition curve. First we consider the phase after a single LD cycle in FD, starting at *t* = 0 and ending at *t* = *T*, where *T* is the period of the LD cycles in FD. The function that gives the phase at the end of the LD cycle is known as the phase transition curve (PTC), *g*(*ϕ*(0)) = *g*(*ϕ*_0_), where

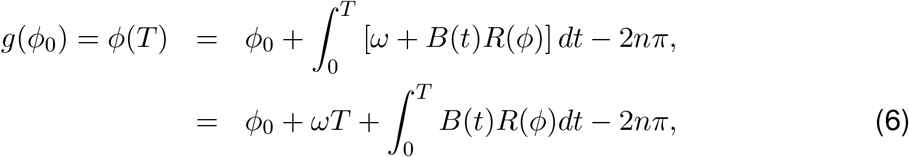

where *n* is the number of times the clock traverses *ϕ* = 0 in the interval from *t* = 0 to *t* = *T* . Equation (6) essentially states that the phase at the end of the cycle is the phase at the beginning of the cycle *ϕ*_0_, plus the phase change due to the intrinsic angular velocity of the clock, *ωT*, plus a phase change due to the effect of the zeitgeber, namely

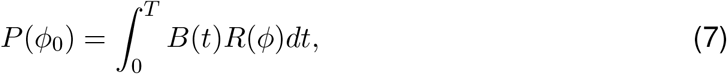

where *P* (*ϕ*_0_) is known as the phase response curve (PRC). Subtracting 2*nπ* ensures that the value of *g*(*ϕ*_0_) remains within the interval [0, 2*π*). The PTC and hence the PRC may be evaluated for any *ϕ*_0_ *∈* [0, 2*π*) by integrating the differential equation in (1).

More generally, defining *ϕ*_*k*_ = *ϕ*(*kT*), then using equations (6) and (7), the phase after each LD cycle is given by the following circle map:

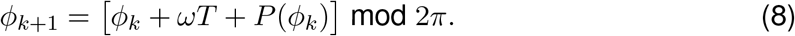

In other words, starting at *ϕ*_0_, after one LD cycle the phase will be *ϕ*_1_, after the next LD cycle the phase will be *ϕ*_2_, etc. This sequence of phases can be visualised by plotting the phase transition curve and constructing the ‘cobweb’ diagram, see Fig. 3 for two examples.

**Figure 3:**
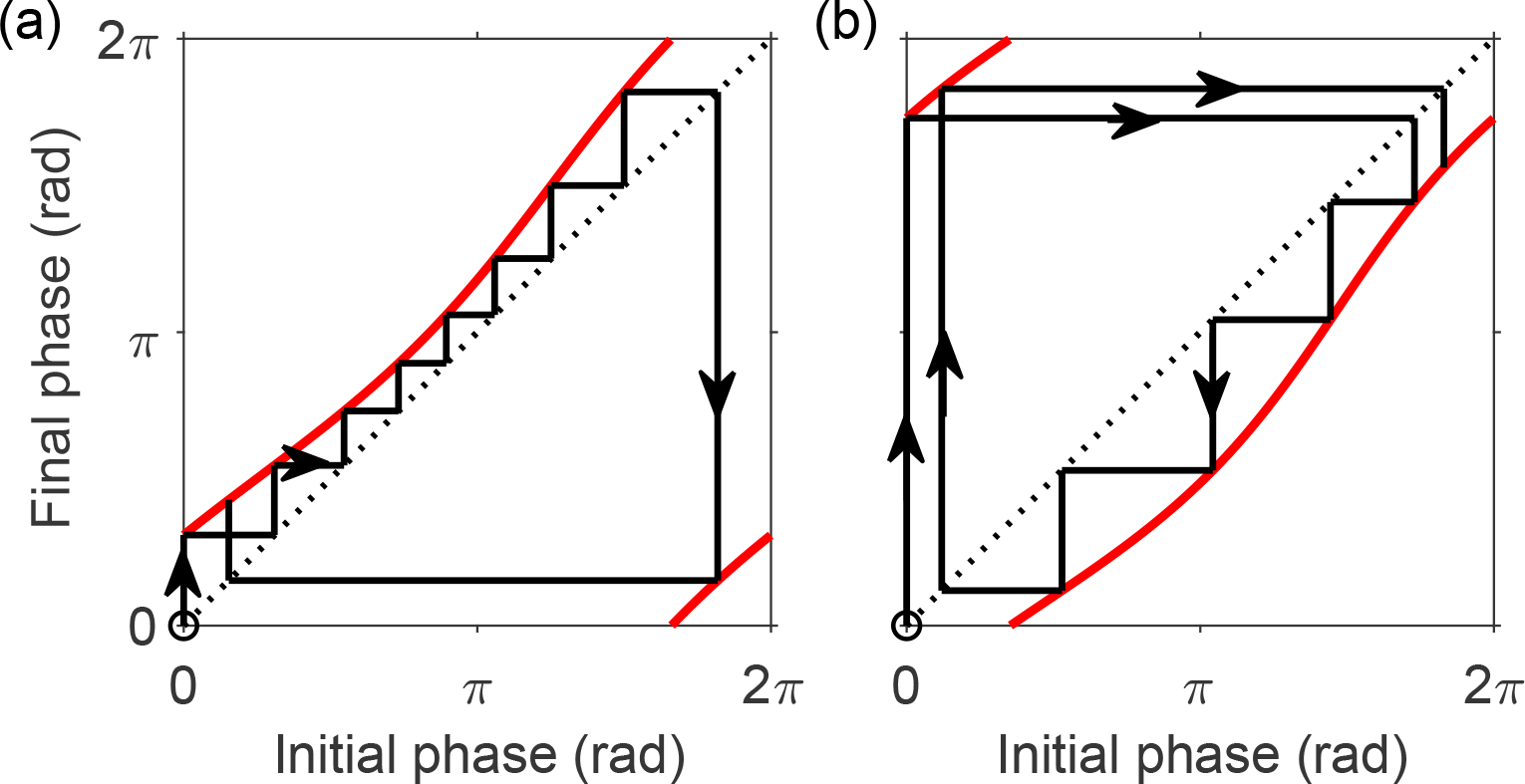
The phase transition curve as a one dimensional map. Cobweb diagrams illustrating the map in (8) in simulations of FD are shown for LD cycles with different periods: (a) *T* =28 h with *M* = 18.7 h i.e. LD cycles with 18.7 h of light and 9.3 h of dark and (b) *T* =20 h with *M* = 13.3 h, i.e. LD cycles with 13.3 h of light and 6.7 h of dark. In each case, the phase transition curve is shown in red. The phase-only model with VRC *R*(*ϕ*) = sin *ϕ* was used with initial phase *ϕ*_0_ = 0.

Having calculated the phase after one LD cycle, we can now calculate the phase after *N* LD cycles as follows. From equation (8),

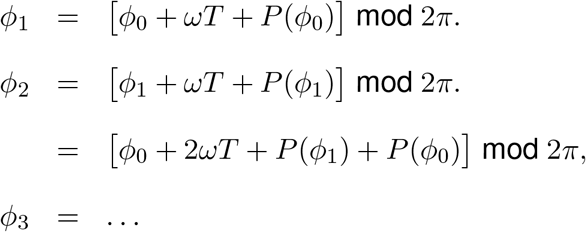

So after *N* cycles,

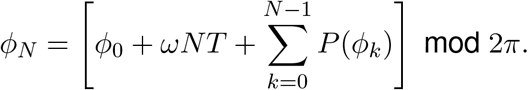

Hence the total change in phase Δ*ϕ*_Tot_ over the course of the FD protocol can be expressed as

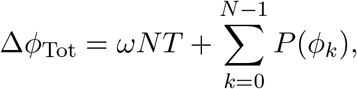

where the first term on the right hand side describes the total change in phase due to the natural angular velocity of the clock and the second term describes the cumulative phase response to successive LD cycles. The mean angular velocity is then:

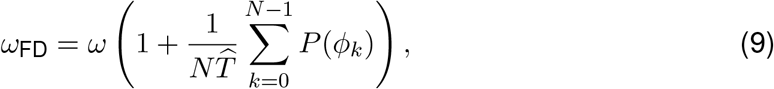

where 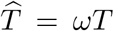. In the next section, we derive an approximate expression for *P* (*ϕ*_*k*_), which leads to an approximate expression for *ω*_FD_.

### Approximate expression for the cumulative phase response across a forced desynchrony protocol

In order to calculate the mean angular velocity *ω*_FD_, and hence period, in forced desynchrony an expression for the cumulative phase response given by the sum term on the right hand side of equation (9) is needed. This is a difficult problem in general, but an approximation can be derived by making two reasonable assumptions, namely that the effect of dim LD cycles is small and that the intrinsic circadian period is close to 24 hours.

We first consider the phase response to a single cycle *P* (*ϕ*_0_). Substituting for *B*(*t*) in equation (7) using equation (2) gives

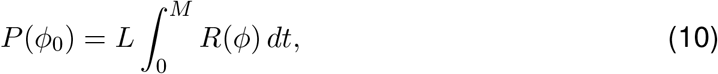

where *M* is the length of the light period and *L* is the magnitude of the stimulus. Using equation (1) to change the variable of integration in equation (10) from *t* to *ϕ* gives an implicit relation between the PRC and the VRC:

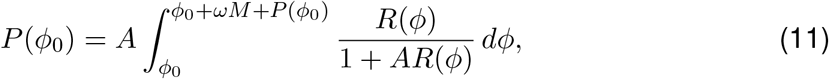

where *A* = *L/ω*. We assume that *A* = *ϵ ≪* 1, that is, the effect of dim light on the angular velocity of the clock is small compared to the intrinsic angular velocity. Then, Taylor expanding the integrand near *ϵ* = 0 and using the fact that *P* (*ϕ*_0_) is *O*(*ϵ*), equation (11) gives

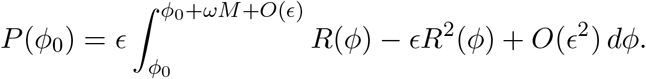

Thus,

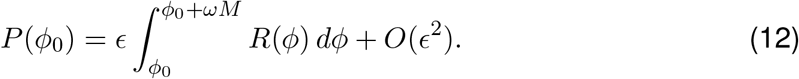

Using the further assumption that the intrinsic period *τ* = 2*π/ω* is close to 24 h, that is *τ* = *T*_solar_(1 + *δ*), where *T*_solar_ = 24 h and |*δ*| *≪* 1, (for example, when *τ* = 24.45 h, *δ*=0.019), in the Supplementary Material we show

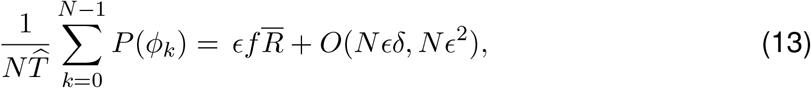

where *f* = *M/T* is the fraction of the LD cycle that is the photoperiod.

### Approximate expression for the period measured in forced desynchrony protocols

Finally, combining equation (13) with (9) leads to an expression for the angular velocity in FD,

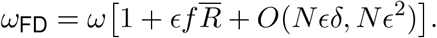

If the *O*(*Nϵδ, Nϵ*^2^) terms are negligible, which means that the stimulus induced by the dim LD cycles is sufficiently small (small *ϵ*) and the intrinsic circadian period is sufficiently close to 24 hours (small *δ*), then

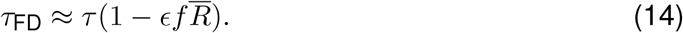

### Implications and accuracy of the approximate expression for τ_FD_

#### Implications of the approximate expression for τ_FD_

Equation (14) states that, to lowest order, the observed circadian period in FD, *τ*_FD_ will be the same as the intrinsic circadian period *τ* . However, unless the VRC has a mean of zero 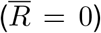 so that it has equal size advance and delay regions, there will be small correction terms. The magnitude of these correction terms will be proportional to the magnitude of the stimulus produced by dim LD cycles *ϵ*, the degree of asymmetry 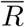 in the VRC and the fraction of the time that the lights are on *f* .

So, if the advance region is larger than the delay region 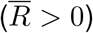 then FD underestimates the intrinsic period. Whereas if the advance region is smaller than the delay region 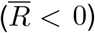 then FD overestimates the intrinsic period. For illustration, equation (14) is plotted in Fig. 4 for three different values of the asymmetry parameter, 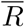, and two different values of the intrinsic circadian period (*τ* =24.15 h and *τ* =24.45 h corresponding to *δ*= 0.006 and *δ*= 0.019 respectively) as a function of the stimulus strength *ϵ*. Hence, as shown in Fig. 4 panel (b), the discrepancy between the period of 24.45 h found in the blind and of 24.15 h found in sighted people in FD protocols in which the lights are on for two-thirds of the time (*f* =2/3), could, for example, be explained by an asymmetry parameter of 0.49 with a stimulus strength of 0.038. Any combination 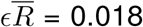 can explain the discrepancy.

**Figure 4:**
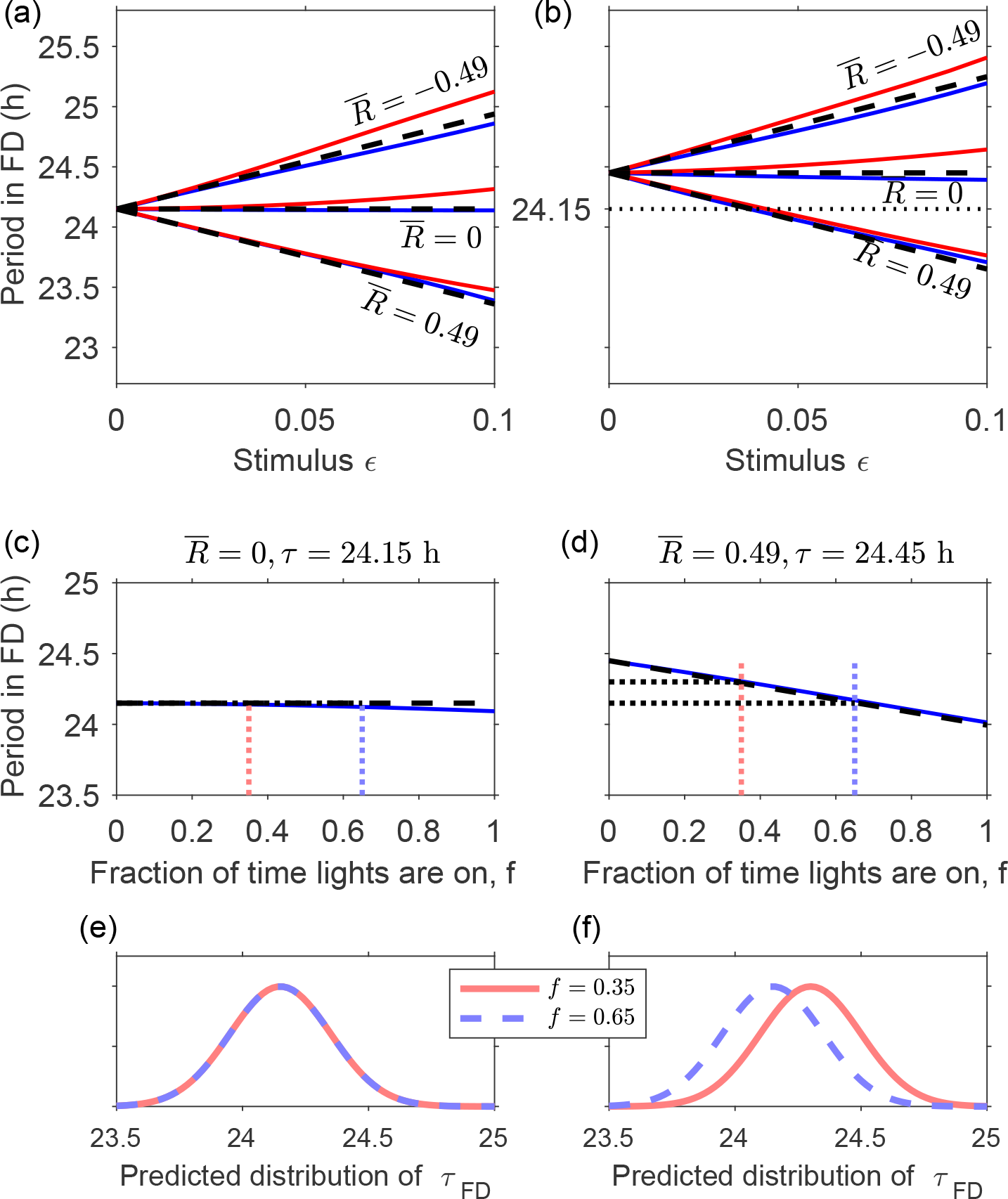
Effect of zeitgeber strength, VRC asymmetry and duration of light period on the observed circadian period in forced desynchrony (FD): predictions using a phase-only model. Upper panels show the period in FD predicted by equation (14) (dashed lines) for a protocol in which the lights are on for two-thirds of the time (*f* =2/3). For comparison, simulations of a 12 LD-cycle FD protocol for *T* =28 h light-dark (LD) cycles (red) and *T* =20 h LD (blue) are shown. Results are shown for three values of the asymmetry parameter 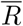 and this for both *τ* =24.15 h (panel (a)), and *τ* =24.45 h (panel (b)). For the simulations, the angular velocity response curve *R*(*ϕ*) = *R*_1_(*ϕ*) = *c* + sin *ϕ* was used, where 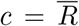, with *ϕ*_0_ = 0. The lower panels show the effect on observed period of the fraction of time that lights are on for 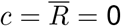(panel(c)) and 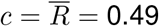 (panel (d)). The light stimulus parameter *ϵ* = 0.038 in both cases. Panels (e) and (f) then show the predicted population distribution of periods measured in FD for a symmetric and an asymmetric VRC respectively.

Here we have chosen to use a stimulus strength of 0.038 because of the fitting discussed below in the ‘Parameter estimation’ section.

Furthermore, equation (14), predicts that the distribution of circadian periods measured in FD will be similar to the distribution of intrinsic circadian periods. For example, if *τ* = 24.45 h, *ϵ* = 0.038 and 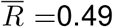, then the standard deviation of *τ*_FD_ will be smaller, but negligibly so, than the standard deviation of *τ* (approximately 2% smaller).

Equation (14) also suggests possible approaches for assessing if there is asymmetry in the VRC. Specifically, since the size of the deviation from the intrinsic period is dependent on the fraction of time in which lights are on, *f*, our analysis predicts that if there is asymmetry in the VRC sufficient to explain the difference between the observed periods in blind and sighted people, then the period in dim 7:13 LD cycles (*f* =0.35), should be approximately 0.14 h longer than the period in dim 13:7 LD cycles (where *f* =0.65). Meanwhile, if the assumption that the VRC is symmetric is valid, there should be no difference between the period in dim 7:13 LD cycles and dim 13:7 LD cycles, see Fig. 4, panels (c)-(f).

### Accuracy of the analytical expression for τFD

In order to illustrate the accuracy of our analytical expression, we compare it to simulations of FD using a simple clock model, see Fig. 4. Equation (14) holds for a general VRC with mean value 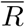. However, for simulations and fitting to data a form for the VRC must be specified. Here, we have taken the lowest order truncation of the Fourier series given in equation (15), namely,

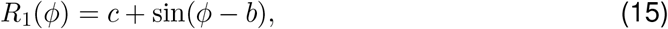

where the VRC to light has been scaled in such a way that the coefficient of the sine term is unity, see Fig. 2. For this specific form for the VRC, 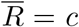. Results are shown for two different values of the cycle length (20 h and 28 h).

Equation (14) assumes that *ϵ* and *δ*are small. Fig. 4 shows that, as is to be expected in an asymptotic analysis, the simulated circadian period deviates from the approximate analytical formula as *ϵ* increases. The deviation is bigger when *δ*is bigger i.e. the deviation from the approximate formula is greater in the right hand panel (*δ*= 0.0208) than in the left (*δ*= 0.00625). Interestingly, the deviation is greater when the period of the LD cycles is *T* = 28 h (red lines) than when *T* = 20 h (blue lines), and this holds regardless of whether the delay region is bigger than the advance region (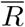 positive) or vice-versa (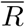 negative).

The deviations of the simulated results from the approximate solution can be explained qualitatively. There are two distinct effects on the observed period as *ϵ* increases. First, as *ϵ* increases, the velocity of the clock is increasingly phase-dependent and, in general, the exact phase response given in equation (10) tends to be smaller than the first-order approximation in equation (12). This effect acts to lengthen the simulated period in FD as compared with the approximate period in equation (14).

Second, as *ϵ* increases, the change in phase from the start of one LD cycle to the next is increasingly non-uniform, so-called relative coordination. The effect of relative coordination on the simulated period depends on the period *T* of the LD cycles. In the left hand panel of Fig. 3 and also in equation (S13) in the Supplementary Material we see that when *T* = 28 h, the phase of the clock advances after each LD cycle, and the advance is smallest when *P* (*ϕ*_*k*_) is minimal. As a result, the clock becomes ‘trapped’ at phases where the PRC is minimal, and relative coordination acts to lengthen the simulated period. Meanwhile, in the right hand panel of Fig. 3, an in equation (S14), we see that when *T* = 20 h, the phase of the clock retreats after each LD cycle, and the retreat is smallest when *P* (*ϕ*_*k*_) is maximal and relative coordination acts to shorten the simulated period.

When the period of the LD cycles in FD is *T* = 20 h, the effect of increasing *ϵ* on the phase-dependent velocity of the clock is balanced to some extent by the effect of relative coordination. Therefore, equation (14) is a better approximation of *τ*_FD_ in *T* = 20 h LD cycles compared to *T* = 28 h LD cycles, especially as *ϵ* increases. Note that all effects are small, so for example, with *f* = 2/3, *ϵ* = 0.038, *c* = 0.49, the analytical expression gives *τ*_FD_ = 24.15 h, the simulations with *T* = 20 h give *τ*_FD_ = 24.14 h, and the simulations with *T* = 28 h give *τ*_FD_ = 24.18 h. It is interesting to note that in Duffy et al. (2011), where results of estimates for *T* = 20 h and *T* = 28 h are given, the mean period observed for *T* = 28 h was longer by 0.04 h than that observed for *T* = 20 h. Since the difference is so small, it was not found to be significant.

### Parameter estimation

Having shown that our analytical formula accurately describes how asymmetry in the VRC and dim LD cycles affect the estimate of intrinsic circadian period in FD in a phase-only model of the circadian pacemaker, the expected next question is whether it is possible to estimate the model parameters from available data. The simple clock model with the sinusoidal VRC contains a total of four parameters, namely the intrinsic period *τ* = 2*π/ω*, the magnitude *L* of the stimulus produced by dim light, the parameter *b* that describes the horizontal shift of the VRC, and the asymmetry parameter *c* (equivalent to 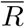) of the VRC. Below, using data from Wright et al. (2001) we estimate the parameters *L* and *b*, although unfortunately we find that it is not possible to estimate *c*.

In Wright et al., four participants were shown to entrain to dim LD cycles with period *T* = 24.0 h. In each participant, their period in a 28-hour FD protocol was also measured. Data for entrained phase were obtained by applying a graph reading application to Fig. 1(b) in Wright’s paper and cross-checked by comparing with results reported in Fig. 2 of Wright et al. (2001) and Fig. 2 of Wright et al. (2005). Wright’s data are reproduced in Fig. 5(a). Circadian phase was measured using DLMO in all cases.

**Figure 5:**
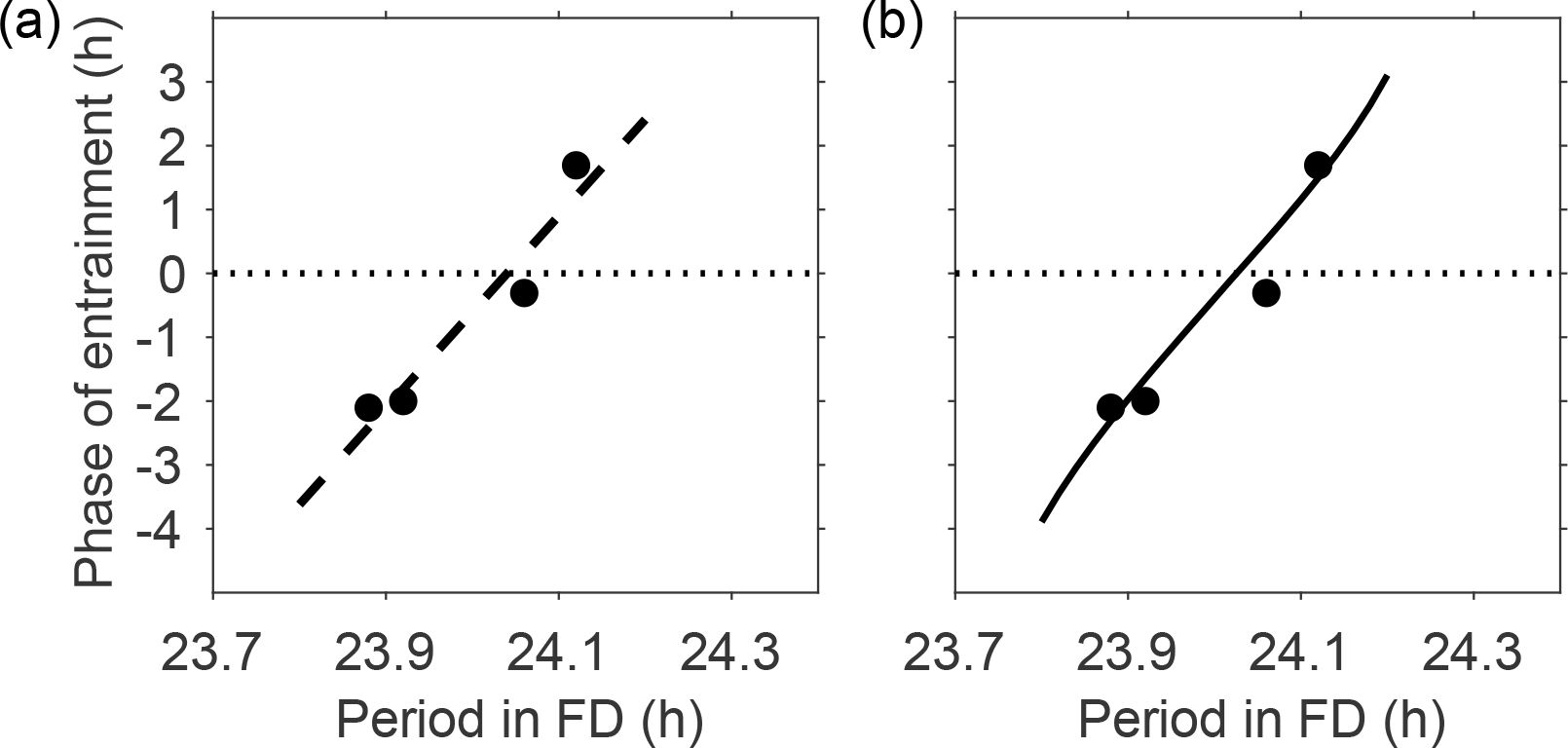
Phase of entrainment to 1.5 lux LD cycles with period T = 24.0 h and photoperiod duration M = 16 h as a function of the period in FD. The phase of entrainment is the timing of DLMO relative to the onset of the dark interval. (a) Data from Wright et al. (2001) and linear regression line fitted to the data. (b) Predictions of the simple clock model with *L* = 0.010 h^*−*1^, *b* =0.63 and *c* = 0 (results are independent of *c*).

The map in equation (8) applies both in FD and during entrainment to LD cycles. Moreover, during entrainment, the clock completes exactly one cycle in each LD cycle, that is

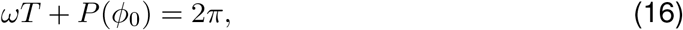

where *ϕ*_0_ is the phase of the clock at the start of the LD cycle. Equation (16) can also be written as

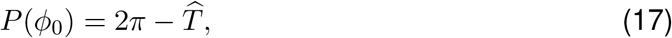

where 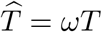, which is known as Pittendrigh’s equation.

In dim LD cycles and making the assumption that the VRC is sinusoidal, equation (12) gives:

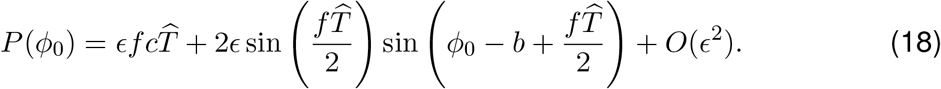

Since *P* (*ϕ*_0_) = *O*(*ϵ*), Pittendrigh’s equation gives 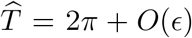. Then, substituting for *P* (*ϕ*_0_) in equation (17) using equation (18) gives

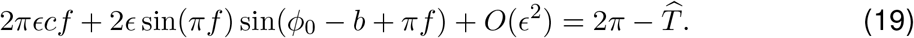

In deriving an explicit expression for the phase of entrainment *Ψ*, we use the arcsin function. Therefore, it is convenient to write equation (19) as

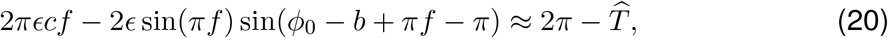

where the argument of the sine function involving *ϕ*_0_ is in the range (*−π/*2, *π/*2).

The phase of entrainment, *Ψ*, reported in Wright et al. (2001) and shown in Fig. 5, is the timing of DLMO relative to the onset of the dark interval of the LD cycles, whereas in (20) phase *ϕ*_0_ is measured from the start of the light interval. Hence *ϕ*_0_ and *Ψ* are related by

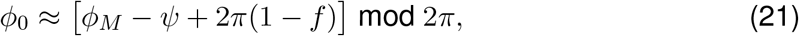

where *ϕ*_*M*_ = 17*π/*12 is the phase of the clock at DLMO, and 2*π*(1 *− f*) is the approximate phase advance between the onset of the dark interval and the onset of the photoperiod.

Substituting equation (21) into equation (20) and rearranging gives an explicit expression for the (stable) phase of entrainment

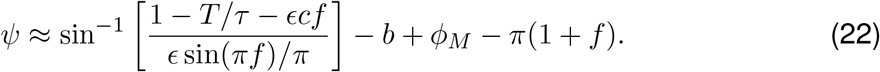

Since *P* (*ϕ*_0_) = *O*(*ϵ*), equation (17) gives *T/τ* = 1 + *O*(*ϵ*), and *T/τ*_FD_ = 1 + *O*(*ϵ*). Then, substituting for *τ* using equation (14) gives *T/τ* = *T/τ*_FD_ *− ϵcf* + *O*(*ϵ*^2^). Thus, equation (22) gives an expression for *Ψ* in terms of *τ*_FD_

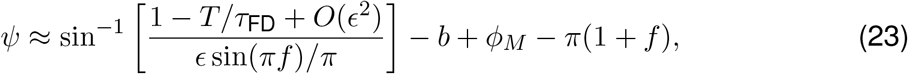

and to the lowest order approximation, the parameter *c* has been eliminated. Next, if we assume *Ψ* is near the middle of the range of entrainment, we can use the small angle approximation to obtain the derivative

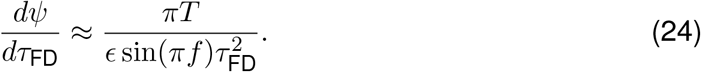

Using measurements of *Ψ, τ*_FD_ and *dΨ/dτ*_FD_, equations (23) and (24) can be solved for *ϵ* and *b*. At the midpoint of the range of Wright’s data, *τ*_FD_ = 24.0 h, and the linear regression line gives

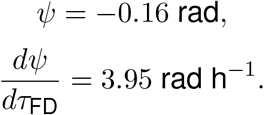

The linear regression line has been superimposed on the data in Fig. 5(a). Using these data and equations (23) and (24), we obtain *ϵ* = 0.038 and *b* = *−*0.63. Since *ϵ* = *L/ω*, for *ϵ* = 0.038 and *τ ≈* 24 h, the fitting suggests that dim light of intensity 1.5 lux produces a stimulus *L ≈* 0.010 h^*−*1^ in our model. Note that the data provide no information on the value of the parameter *c*. The prediction of the phase of entrainment using the simple clock model and the fitted parameters is shown in Fig. 5(b). Since the linear regression line is based on only four data points, the 95% confidence interval for the slope of line is large. Hence, the 95% confidence interval for the parameter *L* based on these data is (0.005 h^*−*1^, 0.085 h^*−*1^). Meanwhile, the 95% confidence interval for *b* is (*−*0.61, *−*0.66).

In another part of Wright’s study, five participants were exposed to dim LD cycles with period *T* = 24.6 h and the period in FD was also measured in these participants. None of the participants entrained to the *T* = 24.6 h LD cycles but their periods were significantly longer than their periods in FD. It is possible to estimate the parameters *ϵ* and *b* in the simple clock model by simulating this part of Wright’s study. This produces similar estimates to those we have obtained using the entrainment data, *ϵ* = 0.034 and *b* = *−*0.84.

### Possible explanation for the difference between the period in FD and the period in DD

Equation (14) relates the period in FD to the intrinsic period. Equation (14) contains the parameter *ϵ*, which represents the strength of the zeitgeber in dim LD cycles, and the parameter 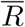, which is the mean value of the VRC. The use of FD to assess the intrinsic period of sighted people is based on the assumption that 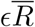 is negligible. In other words, it is assumed that either the VRC to light has equal advance and delay regions 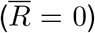, and / or the zeitgeber in FD is too weak to affect the observed period (*ϵ ≈* 0).

However, if 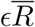 is non-negligible and positive, then this can explain the shorter period in FD compared to the period in DD. For example, using Wright’s study of sighted humans in dim LD cycles, we determined that the stimulus produced by dim light of intensity 1.5 lux is *ϵ* = 0.038. In the FD study reported in Wright et al. (2001), light during waking periods was 1.5 lux and in most other FD protocols it is typically between 5 and 20 lux (Duffy et al., 2011). Together, these suggest that *ϵ* is at least 0.038 during FD protocols.

We expect that light of intensity approximately 15 lux, as frequently used in FD protocols, produces a stimulus of greater magnitude, but without further data we cannot describe how *ϵ* depends on light intensity, although it is likely to be nonlinear. In Kronauer-type models of the circadian pacemaker the light intensity appears raised to the power *p* in the function that describes the action of light on the pacemaker, where *p* is typically taken in the range 0.33 to 0.6 (e.g. see Forger et al., 1999; Jewett et al., 1999; St Hilaire et al., 2007).

## Discussion

The intrinsic period of the central circadian pacemaker is an important parameter that determines the ability of the pacemaker to entrain, and the phase of entrainment, to LD cycles. FD carried out in dim light conditions is considered the gold standard method for measuring circadian period in humans (Dijk and Duffy, 2020; Wang et al., 2023).

We have used a phase-only parametric model to describe the effect of dim LD cycles on the assessment of period in FD. The key result is that the model-predicted period measured in FD, *τ*_FD_ is related to the intrinsic period *τ* by the equation

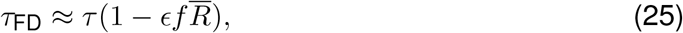

where *ϵ* measures the effect of light, *f* is the fraction of time in each LD cycle for which light is on and 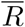 measures the asymmetry of the velocity response curve. The approximation holds when the effect of light is small, as supported by the simulations shown in Fig. 4.

Equation (25) suggests that FD gives a very accurate assessment of intrinsic circadian period provided the VRC to dim light has equal sized advance and delay regions 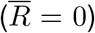 and / or that the stimulus from dim light dark cycles produces a negligible change to the velocity of the clock (*ϵ* small). Using data from entrainment experiments, we have estimated *ϵ* and shown that a positive value of 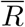, that is, a VRC with a larger advance than delay region, could explain why the mean period of sighted people in FD is shorter than the mean period of free-running blind people.

Our formal estimation of the error between intrinsic circadian period and period as measured in FD also confirms that the design of the protocol means that confounds due to dim light are small. Indeed, we emphasize that this confound is much smaller than that found for sighted individuals in classical free run (see Fig. 1), and we therefore still expect FD to give a much more accurate estimate of intrinsic circadian period than classical free run.

### Period evaluation and after-effects

Several explanations have been offered for differences between the observed circadian period in blind and sighted individuals assessed in FD protocols (e.g. see Czeisler et al., 1999; Lockley et al., 2007; Lewy, 2007). The current dominant view is that observed differences are due to the presence of after-effects i.e. that the period measured in FD in sighted people is a consequence of their prior entrainment to 24 hours. In this scenario, after a sufficiently long time in constant darkness, the period of sighted people would converge to the period observed in the blind. Such long term transients have been observed in nocturnal rodents (Pittendrigh and Daan, 1976).

The simple clock model cannot model after-effects — it responds instantaneously to changes in the light environment, so the mechanism suggested here is fundamentally different. In order to capture after-effects amplitude-phase models (such as the van der Pol oscillator / Kronauer-type models) are required. Indeed, an amplitude-phase model that captures the dependence of measured circadian period on prior light exposure for a diurnal rodent (Bano-Otalora et al., 2021) has been constructed, see Usmani (2022), Chapter 6.

In order to capture after-effects using an amplitude-phase model requires the parameter *μ* to be small (e.g. in Usmani (2022), to fit to the diurnal rodent data, *μ* = 0.02 was taken). However, for humans, the requirement that one 6.7 h pulse of approximately 9500 lux light causes type 1 phase resetting (Khalsa et al., 2003), but pulses on three successive days of 6.7 h of approximately 9500 lux light causes type 0 phase resetting (Khalsa et al., 1997), puts bounds on possible values. Typical values used are in the 0.1-0.25 range, see Supplementary Table 2. A consequence of such large *μ* values is that Kronauer-like models of the human circadian response to light show rapid recovery from perturbations and cannot capture after-effects of the magnitude required to explain the 0.35 h difference in intrinsic period between sighted and blind individuals. At this point, it is not entirely clear how to reconcile both the type 1 and type 0 PRC data and an after-effect interpretation of the period differences between sighted and blind individuals.

### Period evaluation and physiological changes caused by blindness

The only study we have found of sighted people in DD (Wever, 1979, n=5) reported an average intrinsic period consistent with that in the blind. Nevertheless, given the small size of the study, a further explanation is that there are physiological changes in the blind that result in a fundamental change to the circadian system. For example, Yamazaki et al. (2002) show that in hamsters intrinsic circadian period is shorter and more variable following enucleation. Yamazaki et al. suggest a possible explanation is that coupling of retinal clocks with the SCN is an important determinant of the intrinsic period. Hull (2009) also highlights a role for retinal clocks. If there are physiological differences to the circadian system that occur as a result of the lack of light perception, then it may be unrealistic to expect the same mathematical model to describe both blind and sighted people.

### Period evaluation and data selection

A further argument that has been used for the observation of a longer mean intrinsic period in the blind versus sighted people has been that data from studies in the blind have been biased towards those that do not entrain. The accepted value from FD for sighted people is 24.15 h with a standard deviation of 0.20 h (Duffy et al., 2011). In Hull (2009) it was argued that deviations of less than 0.10 h from 24 h could not be detected. If data are normally distributed, then 11% have an intrinsic period less than 23.90 h and 60% have an intrinsic period greater than 24.10 h. The mean of the 60% with a period greater than 24.10 h is 24.28 h. In order to find a mean value of 24.50 h requires selection of the 11% of people with an intrinsic period greater than 24.40 h. For context, in Flynn-Evans et al. (2014), 41 people with no light perception were studied. Of these, 16 (39%) did not entrain and had a mean period of 24.50 h, including one participant with an intrinsic period less than 24 h. Together, these results suggest that bias in data collection cannot explain the magnitude of the difference between sighted and blind individuals.

### Period evaluation and stimulus

We estimated that light of intensity 1.5 lux produces a stimulus *ϵ* of 0.038 based on entrainment data from Wright et al. (2001). This is at least an order of magnitude greater than the value predicted for light of intensity 1.5 lux in the light transduction model of Kronauer (Jewett et al., 1999). Indeed, in order to fit Wright’s data, St. Hilaire et al. (2007) introduced a rest-activity zeitgeber where the rest-activity zeitgeber produced a stimulus of approximately 50 times the magnitude of the light stimulus for light of intensity 1.5 lux. We note that nothing in the derivation of equation (25) explicitly relates to light exposure. Any zeitgeber which can be described in the general form given in equations (1)-(3) will contribute corrections to the intrinsic period of the form 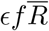 where *ϵ* is the stimulus strength, *f* the fraction of the FD period for which lights are on and 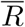 is the mean value of the relevant VRC. This includes the rest-activity zeitgeber introduced in St Hilaire et al. (2007). To first order, contributions will be additive. For example, for two zeitgebers equation (25) becomes

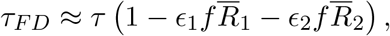

where *ϵ*_*i*_ and 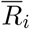 are the stimulus strength and mean value of zeitgeber *i* respectively. However, if rest-activity is the principle zeitgeber of relevance in dim light conditions then one would expect it to have a similar effect in both blind and sighted people.

### Implications and limitations

Mathematical models combined with longitudinal light data collected from people in their natural every day environment, have been suggested as a non-invasive low cost method to estimate circadian phase (Woelders et al., 2017). For day-to-day living the estimates from models are comparable with DLMO. However, models have so far proved less accurate for irregular light-dark schedules as occur during shiftwork (Stone et al., 2019) and when the natural daylength is short (Cheng et al., 2021). Current mathematical models are largely variants of those developed by Kronauer (e.g. Jewett and Kronauer, 1998).

One reason for reduced accuracy in shiftworkers may be that models do not currently adequately capture the response to light levels below 50 lux (e.g. see Fig. 9 in St Hilaire et al., 2007) typical of night-working (Price et al., 2022). Inaccuracy in the modelling of the response to dim light could also explain the reduced accuracy for short natural photoperiods when observed light levels are typically lower (Shochat et al., 2019). One approach to the further development of models is to return to data collected in highly constrained laboratory environments and re-consider whether models adequately capture previous and newly available data, including data on spectral sensitivity of the human circadian pacemaker (e.g. St Hilaire et al., 2022). For example, Usmani (2022) highlights that current models cannot capture circadian phase alignment in both dim and bright light laboratory studies.

Here, we have focussed on period assessment in dim light FD protocols. In dim light, the original Kronauer-type models (e.g. Jewett and Kronauer, 1998) describe a velocity response that is symmetric 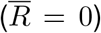. Later versions include additional ‘stimulus modulation’ terms which have the effect of introducing a small amount of asymmetry in the VRC, but substantially less than we propose. Our suggestion that fundamental biological results may be explained by an asymmetric VRC has implications for the design of more accurate mathematical models.

A limitation of our hypothesis is that it depends on the value of the asymmetry parameter 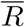. The VRC cannot be measured directly making estimating appropriate values challenging (Taylor et al., 2010). Measured PRC curves to bright light appear approximately symmetric (Khalsa et al., 2003) and may be generated by approximately symmetric VRC’s. However, measuring the PRC in dim light is difficult and it is not clear from current experiments whether the VRC is asymmetric in dim light conditions or not (Revell et al., 2012; St Hilaire et al., 2012). Since the measured phase response in an experimental protocol consists of both a drift due to the intrinsic circadian period and the phase response to light, whether or not PRC’s appear symmetric also depends on the assumed free-running period. We note that others have argued that if there is asymmetry, it is in the opposite direction to the direction we suggest (Khalsa et al., 2003).

We note that our results are consistent with previous simulations using Kronauer-type models in the relevant limits i.e. dim light so that a phase-only model is reasonable, close to symmetric VRC as occurs in Kronauer-type models. Specifically, the simulations of Lok et al. (2022) indicate that a FD protocol with a LD cycle length of 18 h gives a more accurate estimate of the intrinsic circadian period than a 28 h protocol. Stack et al. (2017) simulated an ultradian protocol of 4 h and forced desynchrony protocols of 5 h and 7h and systematically varying light intensity, number of days in the protocol and initial circadian phase. They found that more accurate estimates occurred when light intensity was low and the number of days and length of protocol further facilitated an even distribution of light across circadian phases. In dim light with 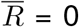, similar to Lok et al., we predict that a 20 h FD protocol gives a more accurate estimate than 28 h. Similar to Stack et al. we find that dimmer light gives more accurate estimates. Where our approach differs is that we have derived an approximate analytical expression which predicts the effect of asymmetry in the VRC on estimates of intrinsic circadian period in dim-light FD protocols.

Finally, validated mathematical models describing the effects of light on the human circadian pacemaker are a prerequisite for understanding the effects of light exposure, which in our society is increasingly dominated by biologically effective man-made light. Novel technologies for monitoring this light exposure longitudinally in people going about their daily lives, combined with validated mathematical models will enable a better prediction of the circadian health consequences of changes in policies related to light exposure and novel light sources.

## Acknowledgements

IMU was supported by the Engineering and Physical Sciences Research Council (grant number: EP/R513350/1).

## Author contributions

Conceptualisation (IM); mathematical analysis (IM, ACS); supervision (ACS, DJD); writing the manuscript (ACS, DJD, IM).

