## Supplementary Material for "Mathematical analysis of light-sensitivity related challenges in assessment of the intrinsic period of the human circadian pacemaker"

5th Oct 2023

### Reduction of typical Kronauer-type models to phase-only models

In the absence of zeitgebers, all Kronauer-type models can be written in the general form

$$\ddot{x} + \tilde{\mu}k(x)\dot{x} + x = 0, \tag{S1}$$

where time has been rescaled, such that $\tilde{t} = \omega t$, where $\omega = 2\pi/\tau_c$. (See Table S1 for the equivalent unscaled form for four popular versions of the model). For small $\tilde{\mu}$, it can be shown that equation (S1) has stable almost sinusoidal solutions $x \approx r(t)\cos(t + \phi(t))$ where $r(t)$ is slowly varying and of $\mathcal{O}(1)$ [1] i.e. a stable limit cycle.

Defining $y = \dot{x}$, and changing to polar coordinates equation (S1) can also be written as

$$\begin{aligned} \dot{x} &= y \\ \dot{y} &= -\tilde{\mu}k(x)y - x. \end{aligned}$$

Changing to polar coordinates using the definitions $x = r\cos\phi, y = r\sin\phi$ gives

$$\begin{aligned} \dot{\phi} &= -1 - \frac{\tilde{\mu}}{2}\sin 2\phi\, k(r\cos\phi) \\ \dot{r} &= -r\tilde{\mu}\sin\phi\, k(r\cos\phi). \end{aligned}$$

For all the Kronauer-type models, close to the limit cycle, $r$ and $k(r\cos\phi)$ are $\mathcal{O}(1)$ and $\tilde{\mu}$ is small. Hence

$$\dot{\phi} \approx -1,$$

i.e. a phase-only oscillator.

The presence of a zeitgeber introduces forcing terms to the van der Pol oscillator equations, but nevertheless, the Kronauer-type models may still be represented in the general form

$$\begin{pmatrix} \dot{x} \\ \dot{y} \end{pmatrix} = \begin{pmatrix} f(x,y) \\ g(x,y) \end{pmatrix} + \mathbf{F}(x,y,t), \tag{S2}$$

where **F** represents the effect of light on the oscillator. The forms of $f(x,y), g(x,y)$ and **F** for four widely used versions are given in Table S1. We note that we have changed variables from definitions in the original papers for ease of comparison between models. For example, for [2], $x_F \equiv -y$ and $y_F \equiv x$.

Again transforming to polar coordinates, we find

$$\dot{\phi} \approx -1 + Bu(\phi, r), \tag{S3}$$

| | Kronauer [3] | Forger [2] | JFK [4] | St Hilaire [5] |
|---|---|---|---|---|
| Oscillator equation | $\left(\frac{12}{\pi}\right)^2 \ddot{x} + \left(\frac{12}{\pi}\right) \mu \left(-1 + 4x^2\right) \dot{x} + \left(\frac{24}{\tau_c}\right)^2 x = 0$ | | $\left(\frac{12}{\pi}\right)^2 \ddot{x} + \left(\frac{12}{\pi}\right) \mu \left(-\frac{1}{3} - 4x^2 + \frac{256}{15}x^6\right) \dot{x} + \left(\frac{24}{\tau_c}\right)^2 x = 0$ | |
| Stiffness parameter | $\mu = 0.13$ | $\mu = 0.23$ | $\mu = 0.13$ | |
| Formulation of oscillator in Cartesian coordinates | $\dot{x} = \left(\frac{\pi}{12}\right) \left[y + \mu \left(x - \frac{4}{3}x^3\right)\right],$ $\dot{y} = -\left(\frac{\pi}{12}\right) \left(\frac{24}{\tau_c}\right)^2 x$ | $\dot{x} = \left(\frac{\pi}{12}\right) y,$ $\dot{y} = \left(\frac{\pi}{12}\right) \left[\mu \left(y - \frac{4}{3}y^3\right) - \left(\frac{24}{f\tau_c}\right)^2 x\right]$ | $\dot{x} = \left(\frac{\pi}{12}\right) \left[y + \mu \left(\frac{1}{3}x + \frac{4}{3}x^3 - \frac{256}{105}x^7\right)\right],$ $\dot{y} = -\left(\frac{\pi}{12}\right) \left(\frac{24}{f\tau_c}\right)^2 x$ | |
| Correction factor to $\tau_c$ | None | $f = 0.99669$ | $f = 0.99729$ | |
| Transduction of light signal $I$ | $B = cI^n$ | $\alpha = \alpha_0 \left(\frac{I}{I_0}\right)^p,$ $\dot{n} = 60\left[\alpha(1-n) - \beta n\right],$ $B = G\alpha(1-n)$ | | $\alpha = \alpha_0 \left(\frac{I}{I_0}\right)^p \left(\frac{I}{I+100}\right),$ $\dot{n} = 60\left[\alpha(1-n) - \beta n\right],$ $B = G\alpha(1-n)$ |
| Transduction parameters | $c = 0.018$ $n = 1/3$ | $\alpha_0 = 0.16$ $I_0 = 9500$ $p = 0.6$ $\beta = 0.013$ $G = 19.875$ | | $\alpha_0 = 0.1$ $I_0 = 9500$ $p = 0.5$ $\beta = 0.007$ $G = 37$ |
| Stimulus modulator | $\widehat{B} = B\left(1 - \frac{1}{3}x\right)$ | $\widehat{B} = B\left(1 - \frac{2}{5}x\right)\left(1 - \frac{2}{5}y\right)$ | | |
| Effect of stimulus in Cartesian coordinates $\mathbf{F}(\mathbf{x})$ | $\left(\frac{\pi}{12}\right) \widehat{B} \begin{pmatrix} 1 \\ qy \end{pmatrix}$ | $\left(\frac{\pi}{12}\right) \widehat{B} \begin{pmatrix} 1 \\ -kx \end{pmatrix}$ | $\left(\frac{\pi}{12}\right) \widehat{B} \begin{pmatrix} 1 \\ qy - kx \end{pmatrix}$ | |
| Drive parameters | $q = 1/3$ | $k = 0.55$ | $q = 1/3, k = 0.55$ | |

Table S1: Comparison of four different versions of Kronauer's model.

where $B$ is the stimulus strength. For dim-light dark cycles, the magnitude of the forcing term is small so $r(t) \approx 1$, hence

$$\dot{\phi} \approx -1 + Bu(\phi, 1), \tag{S4}$$

where and $u(\phi, 1)$ is the equivalent of the velocity response curve in phase only models. For the four models listed in Table S1, the forms of $u(\phi, 1)$ are

$$u(\phi, 1) = \left(1 + \frac{1}{3}\cos\phi\right)\left(\sin\phi + q\sin\phi\cos\phi\right), \tag{S5}$$

for Kronauer's 1998 model [3],

$$u(\phi, 1) = \left(1 + \frac{2}{5}\cos\phi\right)\left(1 - \frac{2}{5}\sin\phi\right)(\sin\phi + q\sin\phi\cos\phi + k\cos^2\phi). \tag{S6}$$

for [4] and [5], and

$$u(\phi, 1) = \left(1 + \frac{2}{5}\cos\phi\right)\left(1 - \frac{2}{5}\sin\phi\right)(\sin\phi + k\cos^2\phi). \tag{S7}$$

for [2]. We note that [5] has an additional non-photic stimulus with velocity response of the form

$$u(\phi, 1) \approx \left[1 + \tanh(10\,\cos\phi)\right]\sin\phi. \tag{S8}$$

Across one cycle, these approximate velocity response curves have mean values of $0, 0.075, 0.0684$ and $0$ respectively. We note that the $\mathcal{O}(\mu)$ terms in equation (S4) are to lowest order odd functions of $\phi$ and independent of $r$ therefore do not contribute to the lowest order terms in the correction to $\tau_{FD}$.

### Evaluating the cumulative phase response

We have,

$$P(\phi_0) = \epsilon\int_{\phi_0}^{\phi_0+\omega M} R(\phi)\,d\phi + O(\epsilon^2). \tag{S9}$$

Substituting for $R(\phi)$ in equation (S9) using the Fourier expansion for the angular velocity response curve, equation (3) gives

$$P(\phi_0) = \epsilon f\overline{R}\widehat{T} + 2\epsilon\sin\left(\frac{f\widehat{T}}{2}\right)\sum_{j=1}^{\infty} a_j\sin\left(j\phi_0 - b_j + \frac{f\widehat{T}}{2}\right) + O(\epsilon^2), \tag{S10}$$

where $f = M/T$ and $\widehat{T} = \omega T$ and we have used the fact that

$$\int_{\phi_0}^{\phi_0+2\gamma} \sin(\phi - b)\, d\phi = 2 \sin\gamma \sin\left(\phi_0 - b + \gamma\right).$$

Similarly, for general $k > 0$, we have

$$P(\phi_k) = \epsilon f \overline{R}\widehat{T} + 2\epsilon \sin\left(\frac{f\widehat{T}}{2}\right) \sum_{j=1}^{\infty} a_j \sin\left[j\phi_k - b_j + \frac{f\widehat{T}}{2}\right] + O(\epsilon^2).$$

Hence,

$$\frac{1}{N\widehat{T}} \sum_{k=0}^{N-1} P(\phi_k) = \epsilon f \overline{R} + \frac{2\epsilon}{N\widehat{T}} \sin\left(\frac{f\widehat{T}}{2}\right) \sum_{j=1}^{\infty} \sum_{k=0}^{N-1} a_j \sin\left[j\phi_k - b_j + \frac{f\widehat{T}}{2}\right] + O(\epsilon^2). \tag{S11}$$

The challenge now is to evaluate the sum over $k$ on the right hand side of equation (S11). We assume that the intrinsic period $\tau = 2\pi/\omega$ is close to $24$ h, that is, $\tau = T_{\text{solar}}(1 + \delta)$, where $T_{\text{solar}} = 24$ h and $|\delta| \ll 1$. (For example, when $\tau = 24.5$ h, $\delta = 0.020$.) Then,

$$\widehat{T} = \omega T = \frac{2\pi T}{\tau} = \frac{2\pi T}{T_{\text{solar}}}\left[1 - \delta + O(\delta^2)\right].$$

Hence, from the map given in equation (8) of the main manuscript,

$$\phi_{k+1} = \left[\phi_k + \frac{2\pi T}{T_{\text{solar}}} + O(2\pi\delta) + O(\epsilon)\right] \text{ mod } 2\pi. \tag{S12}$$

For example, when $T = 28$ h, equation (S12) gives

$$\phi_{k+1} = \left[\phi_k + \frac{\pi}{3} + O(2\pi\delta) + O(\epsilon)\right] \text{mod } 2\pi. \tag{S13}$$

and when $T = 20$ h, equation (S12) gives

$$\phi_{k+1} = \left[\phi_k - \frac{\pi}{3} + O(2\pi\delta) + O(\epsilon)\right] \text{mod } 2\pi. \tag{S14}$$

FD protocols are carried out for a specified number of cycles, where the length of the cycle $T$ and the number of cycles $N$ are frequently chosen so that that $NT$ is an integer number of days. For $T = 28$ h or $T = 20$ h this means that protocols typically last a multiple of six cycles. The significance of equation (S12) is that it shows that for $N$ cycles such that $NT$ is an integer number of days, then the FD protocol will result in $\phi_0, \phi_1, \ldots, \phi_{N-1}$ where the $\phi_k$ are approximately uniformly distributed around the circle.

For evenly spaced $\phi_k = \phi_0 + \frac{2k\pi T}{T_{\text{solar}}}$, a standard calculation can be used as follows. Consider

$$\begin{aligned}\sum_{k=0}^{N-1} \sin\left(j\left(\phi_0 + \frac{2\pi k T}{T_{\text{solar}}}\right) - b_j + \frac{f\widehat{T}}{2}\right) &= \mathrm{Im}\left\{\sum_{k=0}^{N-1} \mathrm{e}^{i\left(j\left(\phi_0 + \frac{2\pi k T}{T_{\text{solar}}}\right) - b_j + \frac{f\widehat{T}}{2}\right)}\right\} \\ &= \mathrm{Im}\left\{\mathrm{e}^{\mathrm{i}\left(\mathrm{j}\phi_0 - \mathrm{b_j} + \frac{\mathrm{f}\widehat{\mathrm{T}}}{2}\right)} \sum_{\mathrm{k}=0}^{\mathrm{N}-1} \mathrm{e}^{\mathrm{ij}\frac{2\pi \mathrm{kT}}{\mathrm{T_{solar}}}}\right\}. \qquad \text{(S15)}\end{aligned}$$

Focussing on the sum term, and letting $\alpha = \frac{2\pi T}{T_{\text{solar}}}$,

$$\sum_{k=0}^{N-1} \mathrm{e}^{ijk\alpha} = 1 + \mathrm{e}^{ij\alpha} + \mathrm{e}^{2ij\alpha} + \ldots + \mathrm{e}^{(N-1)ij\alpha}$$

Hence

$$\sum_{k=0}^{N-1} \mathrm{e}^{ijk\alpha} = \left(\frac{1 - \mathrm{e}^{ijN\alpha}}{1 - \mathrm{e}^{ij\alpha}}\right),$$

and similarly for the real parts of the sum.

FD protocols are chosen so that they are a number of beat cycles, so that $NT$ is an integer multiple of $T_{\text{solar}}$. Hence $\alpha = \frac{2\pi NT}{T_{\text{solar}}}$ is an integer multiple of $2\pi$ and

$$\sum_{k=0}^{N-1} \mathrm{e}^{ijk\alpha} = 0.$$

Hence

$$\sum_{k=0}^{N-1} \sin\left[j\left(\phi_0 + \frac{2\pi k T}{T_{\text{solar}}}\right) - b_j + \frac{f\widehat{T}}{2}\right] = 0,$$

for any $j$. It follows that,

$$\frac{1}{N\widehat{T}} \sum_{k=0}^{N-1} P(\phi_k) = \epsilon f \overline{R} + O(N\epsilon\delta, N\epsilon^2). \qquad \text{(S16)}$$

### Analytical expression for the phase of entrainment for a sinusoidal velocity response curve

Equation (11) is

$$P(\phi_0) = A \int_{\phi_0}^{\phi_0 + \omega M + P(\phi_0)} \frac{R(\phi)}{1 + AR(\phi)}\, d\phi, \qquad \text{(S17)}$$

where $A = L/\omega$.

When

$$R(\phi) = c + \sin(\phi - b),$$

equation (S17) can be integrated. First, the parameter $b$ can be eliminated by a change of variable $\widetilde{\phi} = \phi - b$. Therefore, we proceed to solve

$$\frac{d\phi}{d\widehat{t}} = 1 + A(c + \sin\phi), \tag{S18}$$

where $\widehat{t} = \omega t$. Using the condition in (4), we specify that

$$|A(c + \sin\phi)| < 1. \tag{S19}$$

Therefore,

$$A < \frac{1}{|c + \sin\phi|}. \tag{S20}$$

Note that

$$\max\{|c + \sin\phi|\} = |c| + 1. \tag{S21}$$

Therefore,

$$A < \frac{1}{|c| + 1}, \tag{S22}$$

and we have $A < 1$.

Separating the variables in equation (S18) gives

$$\int_{\phi_0}^{\phi^*} \frac{d\phi}{1 + \widetilde{A}\sin\phi} = \widehat{t}(1 + cA), \tag{S23}$$

where

$$\widetilde{A} = \frac{A}{1 + cA}, \tag{S24}$$

$$\phi_0 = \phi(0), \tag{S25}$$

$$\phi^* = \phi\left(\widehat{t}\right). \tag{S26}$$

Note that $\widetilde{A} < 1$ because of the condition in (S22). Then, using the standard integral

$$\int \frac{d\phi}{1 + \widetilde{A}\sin\phi} = \frac{2}{\sqrt{1 - \widetilde{A}^2}} \arctan\left[\frac{\widetilde{A} + \tan(\phi/2)}{\sqrt{1 - \widetilde{A}^2}}\right] + \text{ constant}, \tag{S27}$$

equation (S23) gives

$$\arctan\left[\frac{\widetilde{A} + \tan(\phi^*/2)}{\sqrt{1 - \widetilde{A}^2}}\right] - \arctan\left[\frac{\widetilde{A} + \tan(\phi_0/2)}{\sqrt{1 - \widetilde{A}^2}}\right] = \frac{\widehat{t}(1 + cA)\sqrt{1 - \widetilde{A}^2}}{2}. \tag{S28}$$

Taking the tangent of equation (S28) and using the angle difference identity

$$\tan(\alpha - \beta) = \frac{\tan\alpha - \tan\beta}{1 + \tan\alpha\tan\beta}, \tag{S29}$$

gives

$$\frac{\left[\widetilde{A} + \tan(\phi^*/2)\right]/F - \left[\widetilde{A} + \tan(\phi_0/2)\right]/F}{1 + \left[\widetilde{A} + \tan(\phi^*/2)\right]\left[\widetilde{A} + \tan(\phi_0/2)\right]/F^2} = C\left(\widehat{t}\right), \tag{S30}$$

where

$$F = \sqrt{1 - \widetilde{A}^2}, \tag{S31}$$

$$C(\widehat{t}) = \tan\left[\frac{F(1 + cA)}{2}\widehat{t}\right]. \tag{S32}$$

Rearranging equation (S30) gives

$$\tan(\phi^*/2) = \frac{C(\widehat{t})/F + \tan(\phi_0/2)\left(1 + \widetilde{A}C(\widehat{t})/F\right)}{1 - \widetilde{A}C(\widehat{t})/F - (C(\widehat{t})/F)\tan(\phi_0/2)}, \tag{S33}$$

and

$$\phi\left(\widehat{t}, \phi_0, A\right) = 2\arctan\left[\frac{C(\widehat{t})/F + \tan(\phi_0/2)\left(1 + \widetilde{A}C(\widehat{t})/F\right)}{1 - \widetilde{A}C(\widehat{t})/F - (C(\widehat{t})/F)\tan(\phi_0/2)}\right] \text{ mod } 2\pi. \tag{S34}$$

During entrainment to laboratory light-dark cycles, Pittendrigh's equation (equation (17) in the main manuscript) is satisfied. Substituting for $P(\phi_0)$ in equation (17) using the phase transition curve in equation (S34) gives

$$\phi\left(f\widehat{T}, \phi_0, A\right) + 2\pi n - \phi_0 - f\widehat{T} = \widehat{\tau} - \widehat{T}, \tag{S35}$$

where $f = M/T$, $A = L/\omega$, and $n$ is the number of times the clock traverses $\phi = 0$ during the photoperiod. Rearranging and taking the tangent of equation (S35), we obtain

$$\tan\left[\frac{\phi_0}{2} - \frac{\widehat{T}(1-f)}{2}\right] = \tan\left[\frac{\phi\left(f\widehat{T}, \phi_0, A\right)}{2}\right]. \tag{S36}$$

Applying the angle difference identity in equation (S29) to the left-hand side of equation (S36) and substituting for $\phi$ using equation (S34) gives

$$\frac{\tan(\phi_0/2) - D}{1 + D\tan(\phi_0/2)} = \frac{C/F + \tan(\phi_0/2)\left(1 + \widetilde{A}C/F\right)}{1 - \widetilde{A}C/F - (C/F)\tan(\phi_0/2)}, \tag{S37}$$

where

$$C = \tan\left[\frac{f\widehat{T}F(1+cA)}{2}\right], \tag{S38}$$

$$D = \tan\left[\frac{\widehat{T}(1-f)}{2}\right]. \tag{S39}$$

Equation (S37) can be rearranged to obtain a quadratic equation in $\tan(\phi_0/2)$:

$$\left[D + \frac{C}{F}\left(1+\widetilde{A}D\right)\right]\tan^2\left(\frac{\phi_0}{2}\right) + \frac{2\widetilde{A}C}{F}\tan\left(\frac{\phi_0}{2}\right) + \left[D + \frac{C}{F}\left(1-\widetilde{A}D\right)\right] = 0. \tag{S40}$$

Since $\tan(\phi_0/2)$ must be real, if there are no real solutions to equation (S40), then entrainment is not possible with the given parameters.

After finding solutions to equation (S40) where they exist, we determine their stability using the condition $P'(\phi_0) < 0$. In simple clock models, the derivative of the phase response curve is

$$P'(\phi_0) = -1 + \frac{F^2(1+C^2)}{C^2\left(1+\widetilde{A}^2\right) + 2\widetilde{A}C(C\sin\phi_0 - F\cos\phi_0) + F^2}. \tag{S41}$$

Once a stable solution for $\phi_0$ has been found, the phase of entrainment can be calculated.
